# From Prefix to Path: Learning Temporally Consistent Biomolecular Dynamics from Limited Initial Data

**DOI:** 10.64898/2026.03.02.709204

**Authors:** Soham Choudhuri, Subinoy Adhikari, Jagannath Mondal

## Abstract

Molecular dynamics (MD) simulations provide detailed insights into biomolecular motion but are often limited by the prohibitive cost of sampling long-timescale behavior. Here, we present a Transformer-based framework that reconstructs temporally continuous dynamical trajectories from only a small fraction of the initial data, directly targeting time-ordered evolution rather than independent ensemble snapshots. Using three systems spanning distinct dynamical regimes (intrinsically disordered *α*-Synuclein, Cytochrome P450 ligand–binding motion, and a synthetic three-well potential), we show that the model learns both local fluctuations and long-range temporal structure. At inference time, the model generates full trajectories autoregressively from an initial prefix as prompt, capturing metastable transitions, basin-to-basin movements, and system-specific dynamical signatures. Free-energy surfaces computed from generated trajectories closely match ground-truth landscapes and, in several cases, we observe enhanced sampling in generated trajectories relative to the trained trajectories—while preserving kinetically meaningful transition patterns. These results demon-strate that Transformer architectures can serve as efficient, system-agnostic tools for time-continuous molecular trajectory prediction, offering a data-driven complement to long MD simulations and enabling accelerated exploration of conformational space.

## Introduction

Molecular dynamics (MD) simulations are indispensable for investigating conformational fluctuations, folding mechanisms, disorder, and functional motions in biomolecular systems.^1^ However, achieving sufficient sampling of long-timescale dynamics remains a persistent challenge due to the substantial computational cost associated with integrating the equations of motion at femtosecond resolution.^2^ This limitation is particularly significant for systems characterized by rugged free-energy landscapes, slow transitions, or broad conformational ensembles, where conventional MD simulation often fails to explore the relevant configurational space within feasible simulation times.^3^

Motivated by this bottleneck, a large fraction of machine-learning approaches in molecular simulation have focused on ensemble generation.^4^ i.e., learning distributions of configurations to improve coverage of conformational space or to propose new states for enhanced sampling. ^3^ While highly successful for accelerating where the system can go, many such methods do not explicitly enforce time continuity: successive generated configurations need not form a dynamically plausible, temporally coherent trajectory consistent with the system’s underlying stochastic evolution. As a result, these approaches^5,6^ can improve equilibrium sampling yet provide limited access to how the system moves across basins, how long it dwells, or which transition pathways dominate—quantities that fundamentally depend on temporally consistent dynamics.

Recent advances in deep learning have demonstrated that neural network, especially those based on the Transformer architecture, are capable of learning temporal patterns from high-dimensional trajectory data.^7,8^ Such models can, in principle, infer the underlying dynamical rules of a system and generate physically plausible continuations of observed trajectories . Leveraging these capabilities to predict full molecular trajectories from only a short initial part of the whole trajectory opens an attractive route toward data-driven acceleration of MD, reducing the need for long simulations while still accessing long-horizon dynamical information. Framing trajectory propagation as a sequence-to-sequence prediction problem therefore offers a direct route to restore time continuity in data-driven acceleration: instead of generating independent samples from an ensemble, the model is trained to extend partial trajectories into long-horizon dynamical realizations.

In this work, we investigate whether a encoder–decoder Transformer can learn to reconstruct complete molecular trajectories from limited initial inputs (1–50% of the trajectory). We evaluate this framework across four distinct systems representing different classes of molecular motion: (i) a synthetic three-well potential system with controlled multi-basin dynamics (1% prefix), (ii) a functional ligand-binding distance coordinate from Cytochrome P450 (full downsampled trajectory used for training, 10% for inference), and (iii) intrinsically disordered protein *α*-Synuclein (50% prefix). These systems collectively span structured folding landscapes, disordered ensembles, metastable functional transitions, and analytically tractable toy models, enabling systematic assessment of generalization across diverse dynamical regimes.

Our results show that the Transformer model can successfully generate long-range trajectory continuations consistent with the underlying physical behavior of each system. Notably, the generated trajectories not only reproduce local fluctuations but also exhibit transitions between free-energy basins, effectively enhancing sampling relative to trained trajectory . Importantly, because the model outputs temporally ordered sequences rather than independent configurations, it provides access to time-resolved dynamical information (e.g., basin dwell patterns and transition occurrences) while simultaneously improving exploration relative to the training trajectories. These results support the view that sequence models can complement MD by providing fast, data-driven trajectory extrapolation that retains time continuity and can aid downstream analyses such as landscape exploration and free-energy estimation.^3^

Overall, this study highlights Transformer-based encoder–decoder architectures as system-agnostic models for temporally coherent molecular trajectory generation, offering a practical route toward machine-learned enhanced sampling with explicit dynamical continuity across biomolecular systems.

## Results

We propose an encoder-decoder Transformer architecture to model and extrapolate molecular dynamics (MD) trajectories from a limited initial segment of the trajectory (figure 1). The model is evaluated on three systems spanning diverse dynamical regimes: the intrinsically disordered protein *α*-Synuclein, a ligand–binding reaction-coordinate trajectory from Cytochrome P450, and a synthetic three-well potential system. Except for the Cytochrome P450 case, trajectories are represented either in a latent two-dimensional space (*z*_1_, *z*_2_) or as a one-dimensional reaction coordinate. The objective is to predict long-time dynamical evolution given only a short prefix of the trajectory, thereby learning the underlying kinetics and transition structure directly from simulation data. Crucially, because the output is a time-ordered continuation conditioned on the prefix, the evaluation emphasizes temporal continuity and kinetic fidelity, not only agreement with marginal (ensemble) distributions.

**Figure 1.**
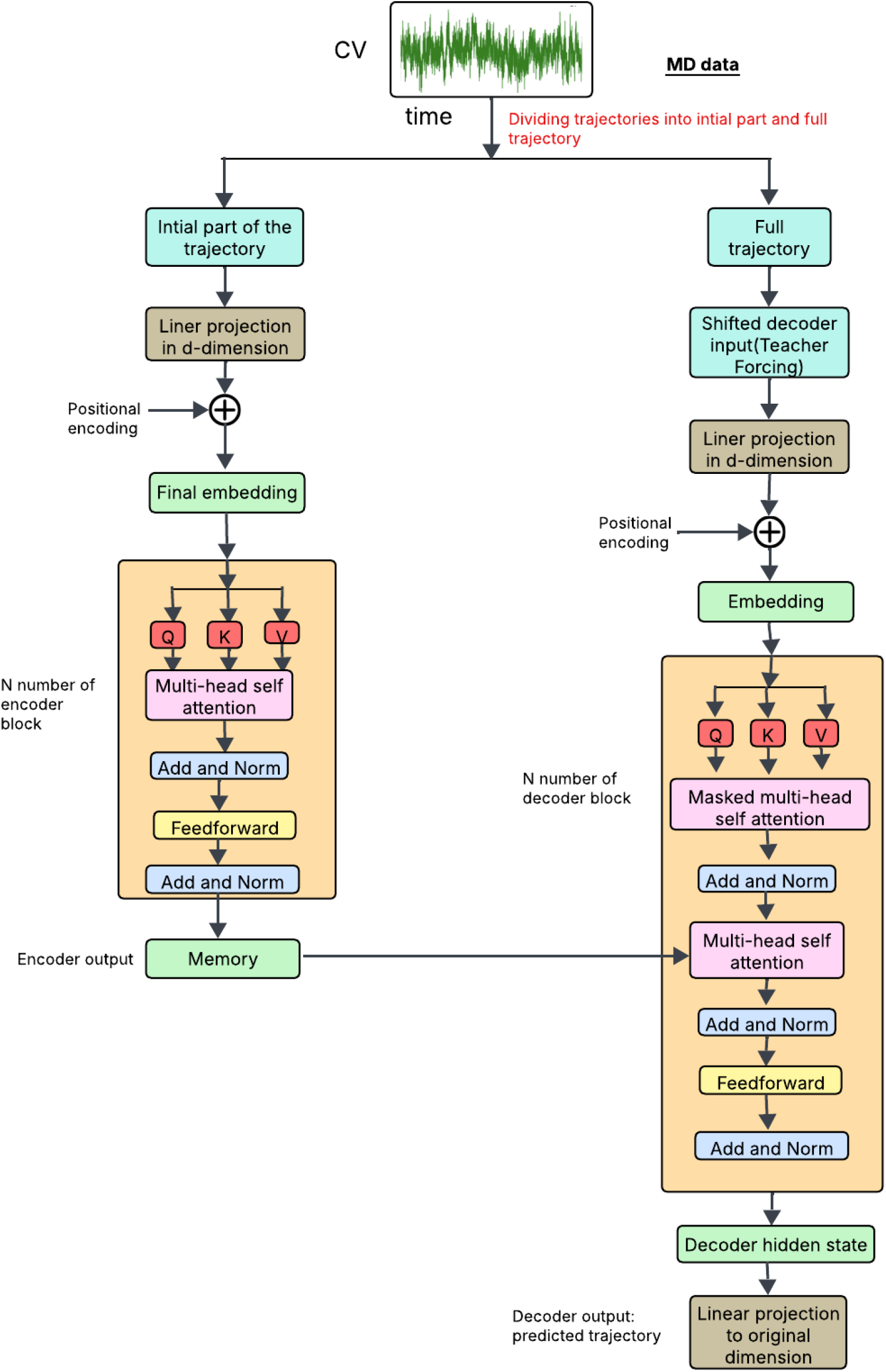
Schematic of the encoder–decoder Transformer for time-continuous trajectory re-construction. The input trajectory is projected into a *d*_model_ embedding and combined with sinusoidal positional encodings to preserve temporal order. The Encoder processes these inputs into a contextual memory representation **M**, capturing long-range dynamical dependencies. The Decoder then uses cross-attention to **M** to autoregressively predict subsequent frames, explicitly enforcing time continuity and enabling the reconstruction of long-timescale molecular transitions from limited initial data.

### Encoder Architecture

Given an initial trajectory prefix of length *S*, denoted by **x**_1:*S*_ *∈ R*^*S×d*^, where *d* is the dimensionality of the input coordinates, the encoder first maps each time step into a *d*_model_-dimensional embedding space using a learnable linear projection,

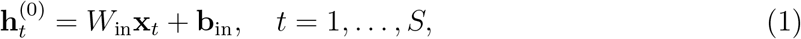

where 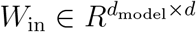 and 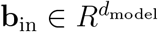 . To preserve temporal ordering, sinusoidal positional encodings **p**_*t*_ are added to the embeddings, yielding

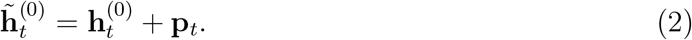

The resulting sequence is processed by a stack of *L*_enc_ Transformer encoder layers, each consisting of a multi-head self-attention module followed by a position-wise feed-forward network, with residual connections and layer normalization applied throughout. For each layer, the self-attention operation is defined as

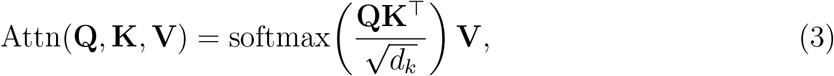

where **Q, K**, and **V** are learned linear projections of the layer input, and *d*_*k*_ is the key dimensionality. This mechanism enables each time step to attend to all others within the prefix, capturing short-time correlations and collective molecular motions. The final encoder output is a contextual memory tensor

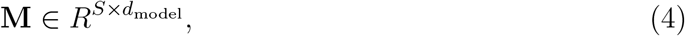

which compactly represents the initial molecular configuration and its early-time dynamics.

### Decoder Architecture and Autoregressive Prediction

During training, the decoder receives a time-shifted version of the ground-truth trajectory **y**_1:*T*_ under teacher forcing. Each decoder layer comprises three submodules: masked multi-head self-attention, encoder–decoder (cross) attention, and a position-wise feed-forward network. The masked self-attention enforces causality by preventing access to future time steps, ensuring that predictions at time *t* depend only on **y**_1:*t−*1_.

Following masked self-attention, cross-attention allows the decoder hidden states to attend to the encoder memory **M**,

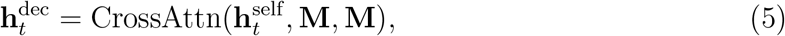

thereby conditioning long-horizon predictions on the encoded initial dynamics. The decoder defines an autoregressive mapping

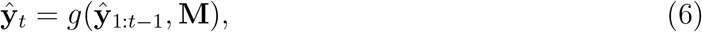

where *g*(·) denotes the composition of masked self-attention, cross-attention, and feed-forward transformations across all decoder layers. The final decoder hidden state is projected back to coordinate space via

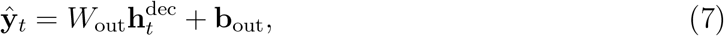

With 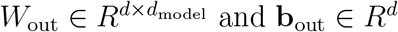.

We have discussed the architecture of our model in detail in the method section and given the mathematical details in the Supplementary.

### Training and Inference

The model is trained by minimizing a mean squared reconstruction loss over the predicted trajectory,

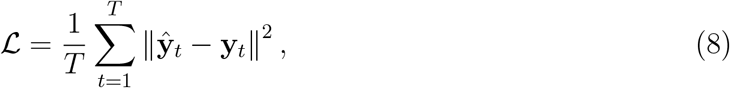

using the Adam optimizer. At inference time, only the short prefix **x**_1:*S*_ is provided to the encoder. The decoder then generates the remaining trajectory sequentially in an autoregressive manner, feeding each newly predicted time step back as input for subsequent predictions. This enables extrapolation of molecular dynamics over long timescales beyond those explicitly observed in the training data. All related codes are given on GitHub: https://github.com/soham2-4/MDgen/tree/main

### Trajectory Reconstruction from Limited Initial Data

Across all four systems, the Transformer model successfully reconstructed long-horizon molecular trajectories from a small fraction of the initial data. Depending on the system, only 1%, 10%, or 50% of the trajectory was provided as input, yet the model generated full-length predictions that preserved the dominant dynamical features. Both visual inspection and quantitative analyses confirmed that the model captured local fluctuations (figure 3), transitions between metastable states, and global dynamical trends, consistent with temporally coherent (time-continuous) evolution, demonstrating its ability to learn effective kinetic representations directly from MD simulations. We have discussed the ITS curves, the CK-test, stationary distributions, FES topologies, and eigenvalue spectra, and generated MSMs in detail in the method section.

### System-Specific Performance

#### Three-Well Potential (Synthetic Multibasin System)

We start by evaluating our Transformer-based method for generating molecular trajectories using a simple but highly informative test case: a two-dimensional Brownian particle moving on a three-well potential energy landscape. This setup is an excellent benchmark, as it includes clearly separated metastable states, predictable transition pathways between them, and dynamics that are close to Markovian—making it straightforward to rigorously compare thermodynamics and kinetics. The reference data come from a long molecular dynamics (MD) simulation on this three-well potential. We split the trajectory into an initial short segment and the whole trajectory. Both represent multivariate sequences tracking the particle’s position over time, specifically the Cartesian coordinates (x(t), y(t)). The Transformer is trained in a conditional reconstruction task: given only the initial segment as input, it learns to generate (reconstruct) the full trajectory. We feed the model raw coordinate values directly to the model. This allows the network to produce continuous trajectories straight in the physical (x, y) configuration space. After training, the model generates new synthetic trajectories through autoregressive sampling, predicting one time step after another. We then subject these generated paths to the exact same thermodynamic and kinetic analyses used on the reference MD data. Visual comparison of representative real and synthetic trajectories (projected onto x(t) and y(t) time series) reveals excellent qualitative match, as illustrated in typical validation figure 2a. The generated paths faithfully reproduce the characteristic stochastic jumps, residence times in basins, and overall temporal structure of the underlying Brownian motion. Thermodynamically, we reconstruct free-energy surfaces (FES) from both the original MD ensemble and the Transformer-generated trajectories Figure 2b and 2c. The synthetic FES closely recovers the three distinct metastable basins, including their correct relative depths, spatial locations, and barrier heights. Crucially, the low-energy connection pathways between basins are preserved with high fidelity. This demonstrates that the model not only matches equilibrium state populations but also learns the global topology of the energy landscape—despite being trained solely on finite-length trajectory fragments. Such agreement strongly suggests the Transformer approximates the stationary distribution driving the dynamics. To assess dynamical fidelity, we discretize configuration space into three states (one per free-energy minimum) and compute state-to-state transition statistics over a range of lag times. Histograms of transition counts—for forward and backward moves between all state pairs—show very close agreement between real and generated data across multiple lag times Figure 2d, 2e and 2f. The decay of transition probabilities with increasing lag follows nearly identical trends in both sets, confirming that the model accurately captures the effective Markovian kinetics and, in particular, the characteristic timescales for barrier crossing. Taken together, the excellent match at the thermodynamic level (via FES reconstruction) and kinetic level (via transition statistics) confirms that attention-based sequence models like the Transformer can reliably reproduce the core physical features of this stochastic dynamical system. A major advantage is that the model outputs full, continuous (x, y) trajectories rather than discrete or coarse-grained samples—allowing direct visualization in configuration space and providing immediate, intuitive evidence of physically realistic behavior. This straightforward low-dimensional validation therefore serves as strong, interpretable evidence supporting the use of Transformer-based approaches for trajectory generation, paving the way for their application to far more complex, high-dimensional biomolecular and soft-matter systems.

**Figure 2.**
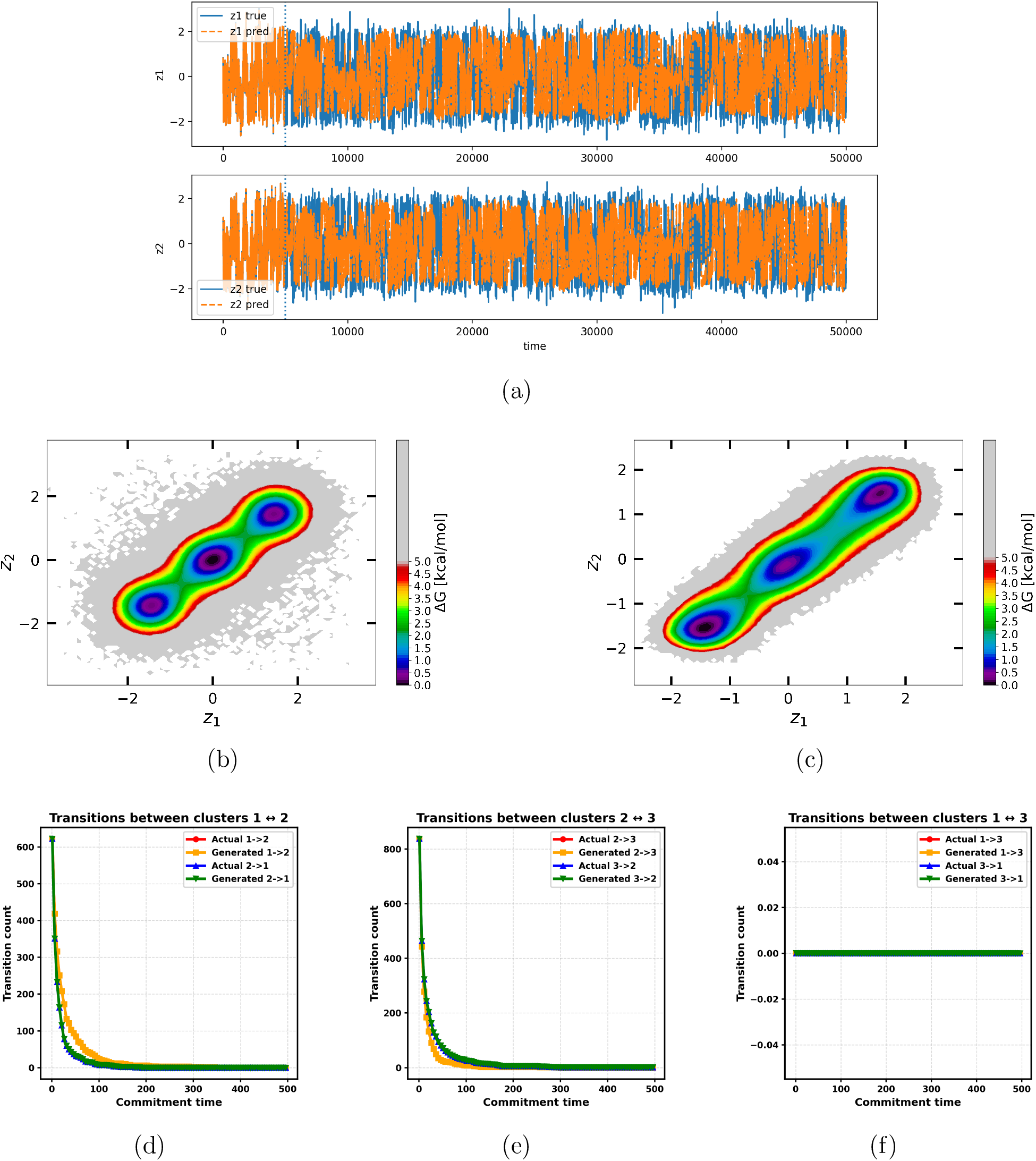
(a) Visual comparison of representative real and synthetic trajectories (projected onto x(t) and y(t) time series) reveals excellent qualitative match(b–c) Free energy surface (FES) of the three-state toy model in the X–Y coordinate space, obtained from (b) the training data and (c) the model-generated data. Transitions between the metastable minima are clearly captured. (d–f) Transition counts as a function of commit time for the three-state toy model. Together, these results demonstrate that the Transformer model successfully learns the contextual relationships among the states and generates trajectories that are both kinetically and thermodynamically consistent.

#### Cytochrome P450 (Ligand–Binding Reaction Coordinate)

We next address one of the most demanding problems in biomolecular modeling: reproducing protein–ligand binding pathways using a Transformer-based generative framework. Binding events, in which a small molecule locates and remains associated with its enzymatic target, are intrinsically rare and typically occur on microsecond-to-millisecond timescales. As a result, conventional unbiased all-atom molecular dynamics simulations are poorly suited for this task, since observing even a single binding event often requires an enormous amount of computation.^9,10^

To demonstrate our approach, we focus on ligand association with cytochrome P450, a well-studied system of substantial biomedical importance (Figure 3a). In our previous work,^11^ obtaining just one successful binding trajectory demanded approximately three to four weeks of uninterrupted GPU computation using GROMACS,^12^ despite it being among the fastest MD engines available. Practical constraints limited us to only three independent simulations, yielding sparse statistics for these infrequent events. This severe computational bottleneck motivates the use of generative models, which aim to amplify a small number of costly simulations into large ensembles of physically realistic trajectories at negligible additional expense.

**Figure 3.**
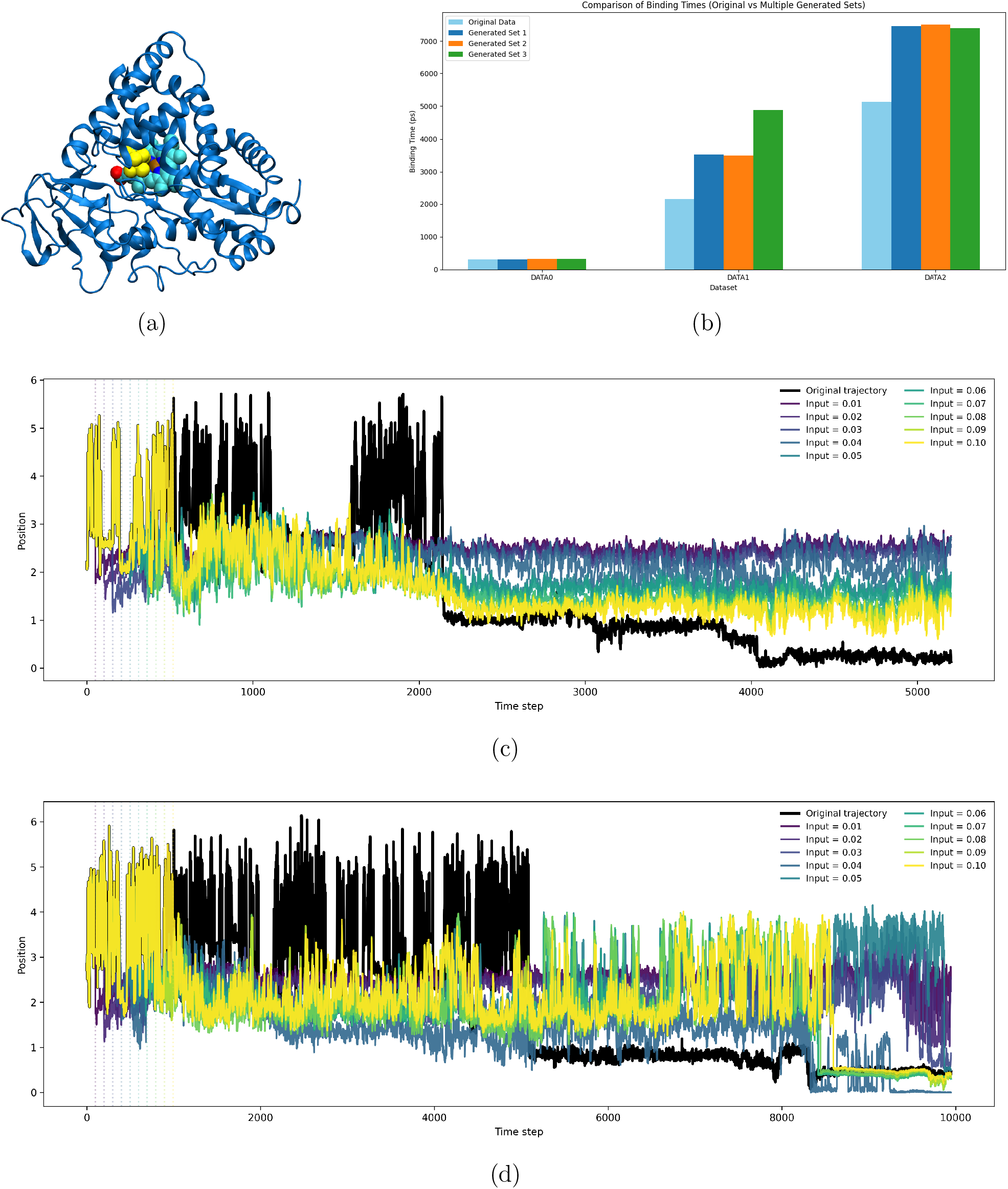
Figures (a): A pictorial representation of Cytochrome P450, Figures (b): Comparison of ligand binding times in the cytochrome P450 system. The bar plot shows close agreement between real MD trajectories and model-generated trajectories, indicating accurate reproduction of P450 binding kinetics. Figure (c), (d): Comparison of real trajectories of protein–ligand distance trajectories for the cytochrome P450 system with generated trajectories from our model using different initial input percentages. We can see that, Input .1 percent means the initial 10% is a good threshold for the generation of trajectories for this system.

The Transformer was trained using these three MD trajectories, represented through a compact yet informative observable: the protein–ligand separation as a function of time. This scalar coordinate effectively summarizes the binding process, characterized by long unbound intervals at large distances, abrupt transitions into close contact, and sustained bound states prior to dissociation.

Each trajectory was partitioned into an initial segment and its corresponding full-length evolution. The model was trained autoregressively to map the early portion of the signal to the complete trajectory by predicting each subsequent distance value conditioned on all previous ones. After training, the Transformer can be seeded with a short initial prompt and used to generate full synthetic distance trajectories autonomously.

We then generated a large ensemble of such trajectories. To assess whether the model captures genuine dynamical behavior rather than merely reproducing average trends, we examined the distribution of binding times—defined as the waiting time to the first successful binding event. Figure 3b compares the distribution obtained from the MD trajectories with that from a large ensemble of Transformer-generated samples. The close agreement between the two is notable, as accurately reproducing rare-event waiting times requires preserving long-range temporal correlations and correct transition kinetics. This result indicates that the attention-based autoregressive model has learned essential aspects of the underlying kinetic landscape, not just static distance statistics. The three MD training traces illustrate the inherent difficulty of the task: long stretches of unbound behavior punctuated by rare, abrupt binding transitions. In contrast, the Transformer-generated trajectories reproduce these features convincingly, displaying extended unbound phases, rapid binding events, and bound-state lifetimes consistent with those observed in the simulations (Figure 3c, 3d).

The computational efficiency gained is striking. Producing the first synthetic trajectory after training required only about few hours, compared to roughly thirty days for a single MD trajectory on the same hardware. This represents an acceleration of several orders of magnitude, transforming the generation of hundreds or thousands of binding trajectories from an impractical endeavor into a routine one.

Overall, these results establish the Transformer as a powerful tool for modeling slow biomolecular association processes. By training on only a few expensive MD trajectories, the model enables rapid generation of large, statistically faithful ensembles of binding pathways. This capability dramatically reduces the computational barrier to studying binding kinetics, mechanistic pathways, and ensemble-level behavior—analyses that would otherwise demand months of large-scale simulation resources. More broadly, this approach offers a practical framework for exploring rare but functionally critical biomolecular events central to enzymatic activity, drug discovery, and related fields.

#### *α*-Synuclein (Intrinsically Disordered Protein)

As a stringent benchmark, we further evaluate our approach on the intrinsically disordered protein *α*-Synuclein,^13,14^ which is characterized by highly heterogeneous conformational dynamics and a rugged free-energy landscape.^15^ To emphasize long-timescale collective behavior, molecular dynamics trajectories are projected onto a two-dimensional latent representation derived from a previously trained *β*-VAE,^16–20^ as illustrated in the Figure in the Supplementary. Latent-space trajectories generated by the Transformer are then directly contrasted with those obtained from MD simulations.

To evaluate whether the generated trajectories preserve correct kinetic behavior, we perform Markov State Model (MSM) analysis. The resulting implied timescale (ITS) plots for the reference MD data and the Transformer-generated data are shown in Figures 4c and 4d, respectively. The close correspondence between these ITS profiles demonstrates that the Transformer successfully reproduces the slow conformational transitions of *α*-Synuclein, even in the presence of a complex and rugged energy landscape. Notably, the ITS curves derived from the generated trajectories exhibit smoother plateaus and faster convergence than those computed from the MD data. This behavior indicates that the model effectively learns the underlying distribution of slow dynamical modes. Moreover, the higher ITS values observed for the Transformer relative to the VAE suggest that the Transformer more accurately captures slower relaxation processes, reflecting an enhanced ability to resolve transitions between kinetically well-separated or metastable states.

**Figure 4.**
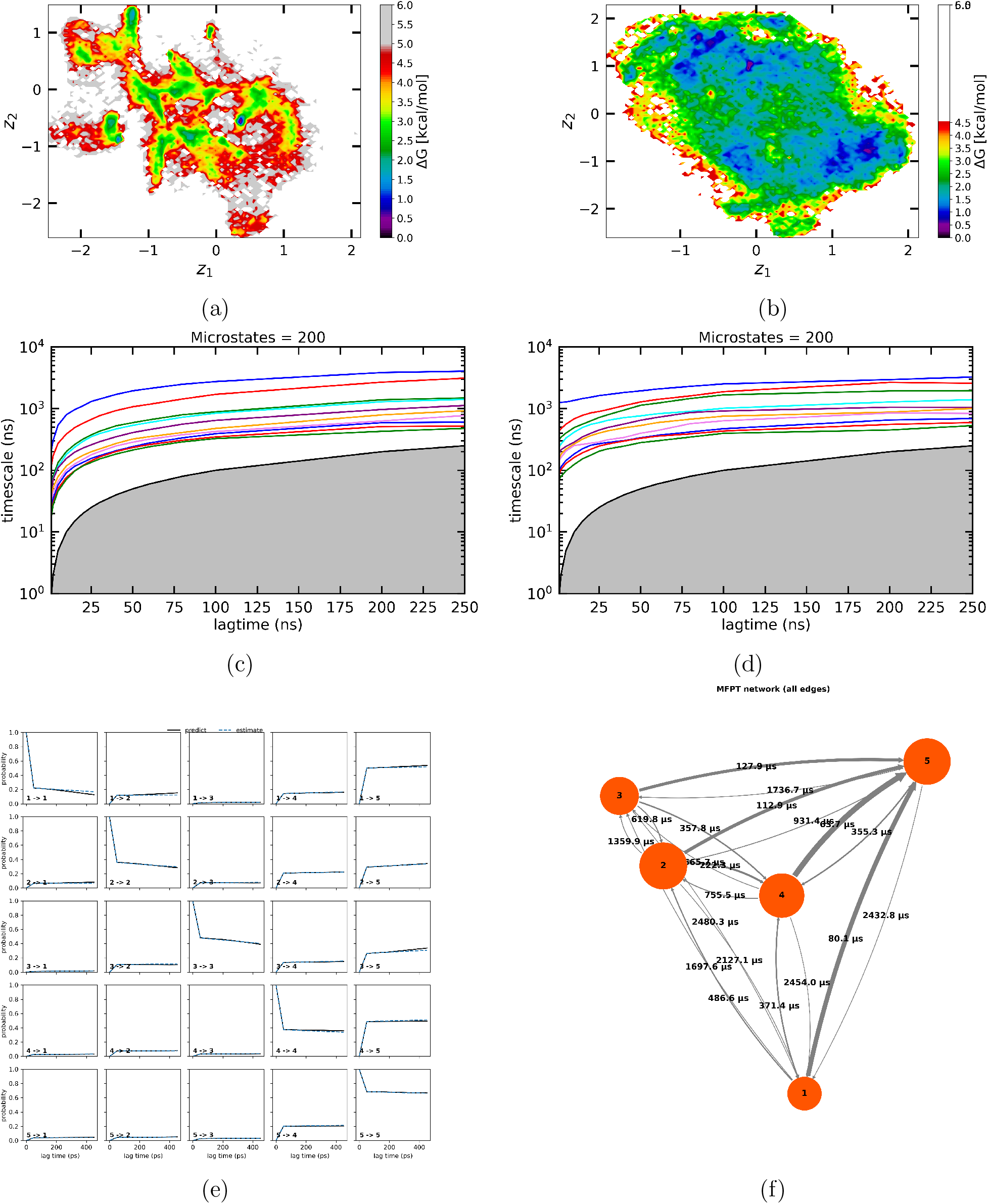
a–b) Free energy surface (FES) of *α*-Synuclein in the latent space, obtained from (a) the training data and (b) the model-generated data. The close similarity between the two landscapes indicates that the model accurately captures the underlying conformational free-energy distribution. Implied timescale (ITS) plots for *α*-Synuclein obtained from (a) the true simulation data and (b) the generated data. (c) CK test for generated data. (d) mean first passage times. (e) Macrostates obtained from PCCA+ along with the cluster centers. (f) Spectral analysis indicating the separatio1n6of spectral states for generated data.

Free-energy surfaces (FES) computed from generated trajectories showed strong qualitative and quantitative agreement with those obtained from ground-truth data. We can see, major dips were recovered with correct topology and depth. For *α*-Synuclein, the generated FES preserved the broad, shallow distribution characteristic of IDPs (figure 4a, 4b). The model-generated trajectories demonstrated *enhanced sampling* relative to the train data, visiting additional metastable states and improving the coverage of the free-energy landscape.

### Generalization Across Dynamical Regimes

Taken together, the results demonstrate that the model generalizes across highly distinct dynamical regimes — from structured folding landscapes to chaotic IDP dynamics and multi-basin synthetic potentials. The architecture successfully integrates information from a short prefix to infer long-range dynamical structure, indicating that Transformer-based sequence models can serve as powerful tools for molecular trajectory prediction and dynamical exploration.

### Model explanation

To gain insight into what the model learns, we conducted a series of interpretability experiments using ligand-binding trajectories of the cytochrome P450 system. After training, the Transformer model is provided with an initial segment of a trajectory and tasked with generating the complete trajectory conditioned on this partial input.

To probe the nature of the learned representation, we systematically scaled the initial trajectory by multiplying or dividing it by a constant factor before feeding it to the model (figure 5b, 5c). Interestingly, despite this rescaling, the model generated full trajectories that preserved the same underlying dynamical structure and semantics, differing only by a corresponding shift in scale. This indicates that the model is not memorizing absolute coordinate values but is instead capturing the relative geometric relationships that govern trajectory evolution.

**Figure 5.**
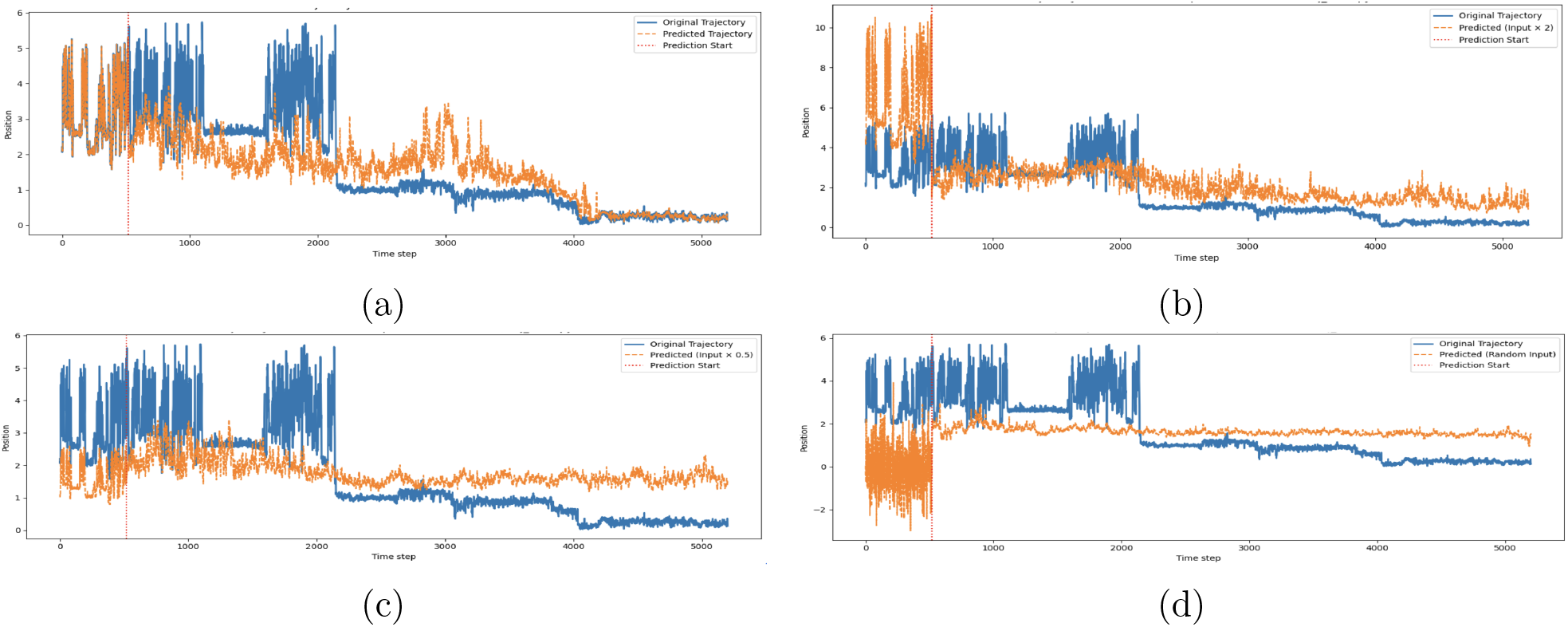
Model interpretability using different initial trajectory inputs. (a) Given the original initial trajectory segment, the Transformer accurately generates the full ligand-binding trajectory. (b and c) When the initial trajectory is uniformly scaled by multiplying by 2 and dividing by 0.5, respectively, the model still produces trajectories that preserve the same underlying geometric and dynamical structure, indicating invariance to absolute scale. (d) In contrast, when a random initial trajectory is provided, the generated trajectory shows no meaningful correspondence with the ground truth. Together, these results demonstrate that the model learns the intrinsic geometric relationship between partial and complete trajectories rather than memorizing absolute coordinate values.

In contrast, when the model was provided with a random initial trajectory (figure 5d) unrelated to the true dynamics, the generated output showed no meaningful correspondence with the ground-truth trajectories (figure 5a) . Together, these observations suggest that the Transformer learns the intrinsic geometric mapping between the initial and full trajectories, rather than relying on superficial numerical patterns.

## Conclusion

In this work, we demonstrate that a Transformer architecture can reliably learn and reproduce time-continuous long-range molecular dynamics from only a small prefix of an input trajectory. Across three systems representing diverse dynamical regimes, intrinsically disordered *α*-Synuclein, Cytochrome P450 ligand-binding motion, and a synthetic three-well potential—the model generated physically meaningful temporally coherent trajectories that capture both short- and long-timescale behaviors. These results highlight the capacity of sequence models to internalize the underlying rules governing molecular motion, even when trained exclusively on latent variables or reaction coordinates.

A key finding is that the model does not simply interpolate between nearby points in the prefix but instead reconstructs the broader dynamical landscape implied by the initial segment. This distinction is central: we aim to preserve temporal ordering and transition statistics, moving beyond ensemble-only generation. In systems such as the three-well potential, the generated trajectories visit multiple metastable basins, indicating that the model has implicitly learned the global topology of the free-energy surface. This enhanced sampling behavior suggests that data-driven generative approaches can complement or accelerate traditional MD by exploring configurational space without requiring long, computationally expensive simulations.

The Transformer model also performs well on challenging systems such as *α*-Synuclein, where the lack of stable structure leads to highly heterogeneous, high-entropy dynamics. The ability to generate non-collapsed, diverse trajectories demonstrates that the architecture can effectively model disorder and remain stable under autoregressive decoding. For functional coordinates such as the P450 ligand-binding distance, the model successfully captures both plateau regions and abrupt transitions, showing that it can learn dynamical features relevant to biochemical function.

Despite these strengths, several limitations remain. Autoregressive prediction can accumulate errors over long times, occasionally leading to drift or loss of fine-grained temporal structure. Integrating probabilistic decoding, physics-informed constraints, diffusion models, or learned energy corrections could further improve stability. Moreover, while this study focuses on latent coordinate trajectories, extending the approach to high-dimensional atomic coordinates is a valuable future direction.

Overall, this work establishes that Transformer-based models offer a promising framework for fast, data-driven molecular trajectory generation. By leveraging a small portion of an observed trajectory to reconstruct the full dynamical evolution, these models provide a potential route toward accelerated sampling, rapid free-energy estimation, and efficient exploration of molecular conformational landscapes. The ability to generalize across fundamentally different systems underscores their suitability as a general tool for modern computational molecular science.

## Methods

### Data Preparation

We evaluated our approach using four molecular systems designed to span diverse dynamical regimes: (i) a three-well potential system, (ii) a ligand–binding reaction-coordinate trajectory from Cytochrome P450,^21^ and (iii) the intrinsically disordered protein (IDP) *α*-Synuclein.^22^ Each trajectory was represented in a latent two-dimensional space.^18^ The two-dimensional representations for *α*-synuclein were obtained from the latent spaces of cosine-annealed and linearly annealed variational autoencoders (VAEs), respectively. The input data for training the VAEs consisted of unbiased molecular dynamics simulation trajectories provided by D. E. Shaw Research, specifically a 73 *μ*s trajectory for α-synuclein. For the three-well toy system, the potential of the system is given as

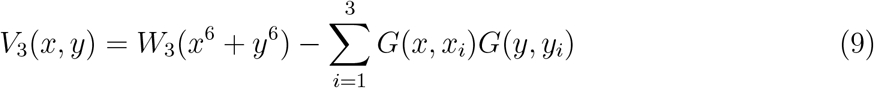

where *G* is a gaussian function as

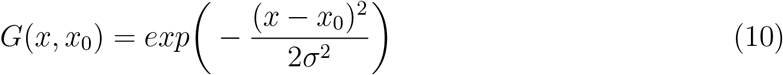

where *x*_0_ and *σ* is the mean and standard deviation of the distribution. In our simulation, we kept *W*_3_ = 0.0001 and *σ* = 0.8. The mean values of Gaussian distribution for the 3-state model are given by *x*_1_ = 0.0, *y*_1_ = 0.0, *x*_2_ = -1.5, *y*_2_ = -1.5, and *x*_3_ = 1.5, *y*_3_ = 1.5. The data was obtained by performing Brownian dynamics simulations by integrating the equation of motion:

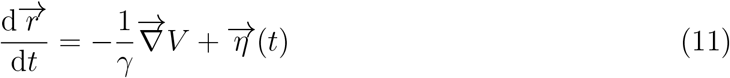

where *γ* is the friction coefficient, *V* is the potential energy, and 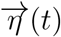 is a random noise, satisfying the fluctuation-dissipation theorem. All simulation times are expressed in units of the Brownian dynamics time scale *τ*_BD_ = *σ*^2^*γ/*(*kT*), where *σ* = 1 denotes the particle diameter. Equation (3) was integrated using the Euler scheme with a time step *δt* = 0.01 *τ*_BD_, and the inverse temperature was fixed at *β* = 1*/*(*kT*) = 9.5. We have divided each trajectories into two parts; initial part of the trajectories and full trajectories. Our aim is to design a generative model which can generate full trajectory from initial part of the trajectory.

Trajectories were uniformly downsampled when necessary and normalized per channel. For each system, only the initial portion of the trajectory (the *prefix*) was provided to the encoder during both training and inference. Prefix lengths were system dependent: 1% (three-well potential), 10% (Cytochrome P450), and 50% (*α*-Synuclein). The full trajectory served as the decoder target during training.

To improve robustness and increase data diversity, additional noise-perturbed trajectories were generated by adding Gaussian noise .0001 to .001. All trajectories were stored at a fixed length *T* for model training.

### Model Architecture

We build an encoder–decoder Transformer architecture to reconstruct full trajectories from a initial part of the trajectory.^8^

#### Transformer Architecture

The initial part of the trajectory **x**_1:*S*_ *∈ R*^*S×*2^ is first projected into a latent space of dimension *d*_model_ using a linear transformation **h**_*t*_ = *W*_in_**x**_*t*_ + **b**_in_, and enriched with sinusoidal positional encodings defined 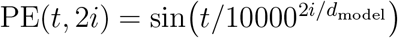 and 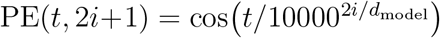, yielding input vectors **z**_*t*_ = **h**_*t*_+PE(*t*). The sequence **z**_1:*S*_ is processed by *L* Transformer encoder layers, each consisting of multi-head self-attention and a positionwise feed-forward network. For each attention head *k*, queries, keys, and values are computed as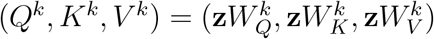, and attention is evaluated as

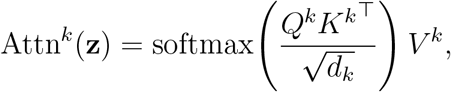

with all heads concatenated and linearly projected to produce the layer output. Stacking *L* layers yields the encoder memory tensor 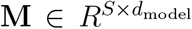. During training, the decoder receives a teacher-forced input 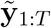 obtained by shifting the ground-truth trajectory by one timestep. Decoder layers apply (i) masked self-attention using a causal mask **C**_*ij*_ = *−∞* for *j > i* to enforce autoregressive prediction,

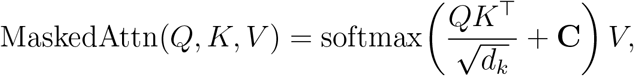

and (ii) cross-attention with queries from the decoder and keys/values from the encoder memory **M**. Autoregressive generation proceeds according to

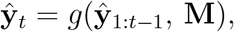

where *g* denotes the composition of masked self-attention, cross-attention, and feed-forward transformations. Finally, each decoder hidden state 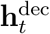 is mapped to the output space via a linear transformation 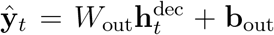, producing the predicted coordinates 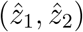and yielding the full autoregressively generated trajectory.

### Training Procedure

The model was trained using the Adam optimizer^23^ with a mean squared error loss:

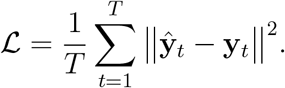

Mini-batches consisted of noise-augmented trajectories. Causal masks were applied to ensure strict autoregressive prediction. Hyperparameters including *d*_model_, number of layers, number of attention heads, and dropout rate were tuned empirically. Model checkpoints were saved periodically.

### Autoregressive Inference

At inference time, only the prefix trajectory **x**_1:*S*_ is available to the model. This prefix is first processed by the encoder to compute a memory tensor **M**, which summarizes the contextual information contained in the observed segment. The decoder is then initialized with a start token and generates the trajectory autoregressively, predicting one timestep at a time using masked self-attention together with cross-attention over **M**. Each newly predicted point is appended to the decoder input and used to condition subsequent predictions, and this process is repeated until the full trajectory of length *T* is generated. This inference procedure enables the model to extrapolate long-range dynamical behavior from minimal initial information.

### Kinetics Analysis via Markov State Modeling

Markov state models (MSMs) provide a rigorous framework for quantifying the long-timescale kinetics of molecular systems, making them essential for validating whether generated trajectories reproduce the true dynamical behavior of the underlying MD simulations.^24–29^ While geometric similarity alone can indicate whether trajectories look physically plausible, only MSM analysis can reveal whether key kinetic properties—such as metastable state structure, transition probabilities, relaxation timescales, and slow dynamical modes—are faithfully captured. By comparing implied timescales, Chapman–Kolmogorov consistency, spectral eigenvalues, and stationary distributions between the training and generated trajectories, MSMs allow us to assess whether the Transformer model has learned not just the shape of the trajectory but also the correct kinetic evolution on the free-energy landscape.^30,31^ Thus, MSM analysis serves as a stringent test of whether the generative model preserves the true physics of molecular motion.

To evaluate whether the Transformer-generated trajectories reproduce the kinetic behavior of the underlying molecular systems, we constructed Markov state models (MSMs) from both the ground-truth MD trajectories and the model-generated trajectories. All analyses were performed using standard MSM workflows, including state discretization, transition probability estimation, and kinetic validation tests using PyEMMA.^32^ All related codes are given on GitHub: https://github.com/soham2-4/MDgen/tree/main

### Construction of the Markov State Model

For the alpha-synuclein, the true and the generated trajectory was discretized into a set of microstates using *k*-means clustering. The optimum number of clusters were chosen based on the VAMP2 score (this needs to be done). We chose 200/500 microstates to ensure a detailed representation of the true and generated trajectory. This was followed by construction of a transition matrix by counting transitions between microstates at a given lag time. To determine an appropriate lag time, the implied timescales were computed over a range of lag times and plotted as a function of lag time. The lag time corresponding to the plateau of the implied timescale curves was selected for constructing the final Markov state model (MSM).

#### Transition Matrix Estimation

Given a chosen lag time *τ*, transition probability matrices were estimated as

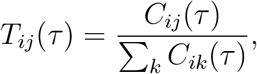

where *C*_*ij*_(*τ*) denotes the number of transitions observed from state *i* to state *j* after lag *τ* . Separate MSMs were built for: (i) the original MD training trajectories and (ii) the Transformer-generated trajectories.

#### Implied Timescales (ITS)

Implied timescales were computed from the eigenvalues *λ*_*k*_ (*τ*) of the transition matrices via

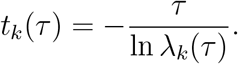

ITS curves were compared between training and generated data to evaluate whether the long-timescale relaxation processes were preserved. Agreement in the slope and plateau regions of ITS curves indicates kinetic consistency.

#### Chapman–Kolmogorov Test

The Chapman–Kolmogorov (CK) test was used to assess the Markovianity and predictive accuracy of both MSMs. Predicted state probabilities at lag *nτ*,

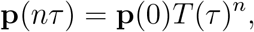

were compared with empirical estimates obtained directly from the trajectories. Consistency between predicted and observed probabilities indicates that the generated trajectories obey the same kinetic evolution as the original MD data.

#### Free-Energy Surface Comparison

Free-energy surfaces (FES) were estimated from the stationary distributions of the MSMs using

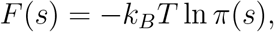

where *π*(*s*) is the stationary probability of state *s*. FES from generated trajectories were compared with those from training data to evaluate whether major basins, barrier heights, and global landscape topology were preserved.

#### Spectral Analysis

The spectral properties of the transition matrices were compared by examining the leading eigenvalues and eigenvectors. Correspondence in the spectral gap and dominant eigenmodes indicates that the generated trajectories reproduce key slow dynamical processes. Differences in higher eigenmodes were used to identify deviations in fast-timescale dynamics.

#### Overall Kinetic Consistency

Across all systems, we compared the ITS curves, CK-test results, stationary distributions, FES topologies, and eigenvalue spectra between ground-truth and generated MSMs. Agreement across these metrics indicates that the Transformer model not only reconstructs structural or geometric properties of trajectories but also captures the underlying kinetics and the global organization of the molecular free-energy landscape.

## Data and code availability

The codes developed in this project is publicly available in Github (https://github.com/soham2-4/MDgen/tree/main)

## Acknowledgement

We acknowledge support of the Department of Atomic Energy, Government of India, under project Identification number: RTI 4007. JM acknowledges the core research grant approved by the Department of Science and Technology (DST) of India (CRG/2023/001426).

## Supporting Information Available

Supplemental information includes a mathematical description of the model.

## References

(1) Karplus, M.; McCammon, J. A. Molecular dynamics simulations of biomolecules. Nature structural biology 2002, 9, 646–652.

(2) Shaw, D. E.; Deneroff, M. M.; Dror, R. O.; Kuskin, J. S.; Larson, R. H.; Salmon, J. K.; Young, C.; Batson, B.; Bowers, K. J.; Chao, J. C.; others Anton, a special-purpose machine for molecular dynamics simulation. Communications of the ACM 2008, 51, 91–97.

(3) Wang, Y.; Ribeiro, J. M. L.; Tiwary, P. Machine learning approaches for analyzing and enhancing molecular dynamics simulations. Current opinion in structural biology 2020, 61, 139–145.

(4) Tiwary, P.; Herron, L.; John, R.; Lee, S.; Sanwal, D.; Wang, R. Generative AI for computational chemistry: A roadmap to predicting emergent phenomena. Proceedings of the National Academy of Sciences of the United States of America 2025, 122, e2415655121.

(5) Noé, F.; Olsson, S.; Köhler, J.; Wu, H. Boltzmann generators: Sampling equilibrium states of many-body systems with deep learning. Science 2019, 365, eaaw1147.

(6) Feng, B.; Zhang, J.; Zhang, X.; Liu, Z.; Li, Y. BioMD: All-atom Generative Model for Biomolecular Dynamics Simulation. arXiv preprint arXiv:2509.02642 2025,

(7) Jing, B.; Stärk, H.; Jaakkola, T.; Berger, B. Generative modeling of molecular dynamics trajectories. Advances in Neural Information Processing Systems 2024, 37, 40534–40564.

(8) Vaswani, A.; Shazeer, N.; Parmar, N.; Uszkoreit, J.; Jones, L.; Gomez, A. N.; Kaiser, L-.; Polosukhin, I. Attention is all you need. Advances in neural information processing systems 2017, 30 .

(9) Shan, Y.; Kim, E. T.; Eastwood, M. P.; Dror, R. O.; Seeliger, M. A.; Shaw, D. E. How Does a Drug Molecule Find Its Target Binding Site? Journal of the American Chemical Society 2011, 133, 9181–9183.

(10) Ahalawat, N.; Mondal, J. An Appraisal of Computer Simulation Approaches in Elucidating Biomolecular Recognition Pathways. The Journal of Physical Chemistry Letters 2020, 12, 633–641.

(11) Ahalawat, N.; Mondal, J. Mapping the Substrate Recognition Pathway in Cytochrome P450. Journal of the American Chemical Society 2018, 140, 17743–17752.

(12) Pronk, S.; Páll, S.; Schulz, R.; Larsson, P.; Bjelkmar, P.; Apostolov, R.; Shirts, M. R.; Smith, J. C.; Kasson, P. M.; Van Der Spoel, D.; others GROMACS 4.5: a highthroughput and highly parallel open source molecular simulation toolkit. Bioinformatics 2013, 29, 845–854.

(13) Goedert, M. Alpha-synuclein and neurodegenerative diseases. Nature Reviews Neuroscience 2001, 2, 492–501.

(14) Butler, B.; Sambo, D.; Khoshbouei, H. Alpha-synuclein modulates dopamine neurotransmission. Journal of chemical neuroanatomy 2017, 83, 41–49.

(15) Chen, J.; Zaer, S.; Drori, P.; Zamel, J.; Joron, K.; Kalisman, N.; Lerner, E.; Dokholyan, N. V. The structural heterogeneity of α-synuclein is governed by several distinct subpopulations with interconversion times slower than milliseconds. Structure 2021, 29, 1048–1064.

(16) Adhikari, S.; Mondal, J. Machine Learning Subtle Conformational Change due to Phosphorylation in Intrinsically Disordered Proteins. The Journal of Physical Chemistry B 2023,

(17) Menon, S.; Adhikari, S.; Mondal, J. An Integrated Machine Learning Approach Delineates an Entropic Expansion Mechanism for the Binding of a Small Molecule to α-Synuclein. eLife 2024, 13, RP97709.

(18) Adhikari, S.; Mondal, J. Elucidating Protein Dynamics through the Optimal Annealing of Variational Autoencoders. Journal of Chemical Theory and Computation 2025, 21, 6367–6379.

(19) Sarkar, D. K.; Adhikari, S.; Surolia, A. Leveraging Deep Learning and MD Simulations to Decipher the Molecular Basis of Attenuated Activity in Glycocin F. bioRxiv 2025, 2025–10.

(20) Be, N.; Adhikari, S.; Mondal, J. Fast Adversarial Generation of Molecular Dynamics Trajectories with Kinetic Fidelity. bioRxiv 2026, 2026–01.

(21) Klepeis, J. L.; Lindorff-Larsen, K.; Dror, R. O.; Shaw, D. E. Long-timescale molecular dynamics simulations of protein structure and function. Current opinion in structural biology 2009, 19, 120–127.

(22) Robustelli, P.; Piana, S.; Shaw, D. E. Developing a molecular dynamics force field for both folded and disordered protein states. Proceedings of the National Academy of Sciences 2018, 115, E4758–E4766.

(23) Adam, K. D. B. J.; others A method for stochastic optimization. arXiv preprint arXiv:1412.6980 2014, 1412.

(24) Prinz, J.-H.; Wu, H.; Sarich, M.; Keller, B.; Senne, M.; Held, M.; Chodera, J. D.; Schütte, C.; Noé, F. Markov models of molecular kinetics: Generation and validation. The Journal of chemical physics 2011, 134 .

(25) Husic, B. E.; Pande, V. S. Markov state models: From an art to a science. Journal of the American Chemical Society 2018, 140, 2386–2396.

(26) Bowman, G. R.; Pande, V. S.; Noé, F. An introduction to Markov state models and their application to long timescale molecular simulation; Springer Science & Business Media, 2013; Vol. 797.

(27) Wu, H.; Paul, F.; Wehmeyer, C.; Noé, F. Multiensemble Markov models of molecular thermodynamics and kinetics. Proceedings of the National Academy of Sciences 2016, 113, E3221–E3230.

(28) Noé, F.; Rosta, E. Markov models of molecular kinetics. The Journal of chemical physics 2019, 151 .

(29) Shukla, D.; Hernández, C. X.; Weber, J. K.; Pande, V. S. Markov state models provide insights into dynamic modulation of protein function. Accounts of chemical research 2015, 48, 414–422.

(30) Chodera, J. D.; Noé, F. Markov state models of biomolecular conformational dynamics. Current opinion in structural biology 2014, 25, 135–144.

(31) Noé, F.; Clementi, C. Kinetic distance and kinetic maps from molecular dynamics simulation. Journal of chemical theory and computation 2015, 11, 5002–5011.

(32) Scherer, M. K.; Trendelkamp-Schroer, B.; Paul, F.; Pérez-Hernández, G.; Hoffmann, M.; Plattner, N.; Wehmeyer, C.; Prinz, J.-H.; Noé, F. PyEMMA 2: A software package for estimation, validation, and analysis of Markov models. Journal of chemical theory and computation 2015, 11, 5525–5542.

