## Supplemental Text for "From Prefix to Path: Learning Temporally Consistent Biomolecular Dynamics from Limited Initial Data"

February 17, 2026

### 1 mathematical details of the model

**Problem formulation** Given an observed prefix $\mathbf{x}_{1:S} \in \mathbb{R}^{S\times 2}$, we model the conditional distribution of a full trajectory $\mathbf{y}_{1:T} \in \mathbb{R}^{T\times 2}$ as

$$p(\mathbf{y}_{1:T} \mid \mathbf{x}_{1:S}) = \prod_{t=1}^{T} p(\mathbf{y}_t \mid \mathbf{y}_{1:t-1}, \mathbf{x}_{1:S}). \tag{1}$$

**Input embedding** Each coordinate $\mathbf{x}_t \in \mathbb{R}^2$ is linearly embedded,

$$\mathbf{h}_t = W_{\text{in}}\mathbf{x}_t + \mathbf{b}_{\text{in}}, \quad W_{\text{in}} \in \mathbb{R}^{d_{\text{model}}\times 2}. \tag{2}$$

Sinusoidal positional encodings are added,

$$\text{PE}(t, 2i) = \sin\left(\frac{t}{10000^{2i/d_{\text{model}}}}\right), \tag{3}$$

$$\text{PE}(t, 2i+1) = \cos\left(\frac{t}{10000^{2i/d_{\text{model}}}}\right), \tag{4}$$

yielding encoder inputs

$$\mathbf{z}_t = \mathbf{h}_t + \text{PE}(t), \qquad \mathbf{Z} = [\mathbf{z}_1, \ldots, \mathbf{z}_S]^\top. \tag{5}$$

**Encoder self-attention** For head $k$,

$$Q^k = \mathbf{Z}W_Q^k, \quad K^k = \mathbf{Z}W_K^k, \quad V^k = \mathbf{Z}W_V^k, \tag{6}$$

with $W_Q^k, W_K^k, W_V^k \in \mathbb{R}^{d_{\text{model}}\times d_k}$, $d_k = d_{\text{model}}/H$. Scaled dot-product attention is

$$\text{Attn}^k(\mathbf{Z}) = \text{softmax}\left(\frac{Q^k(K^k)^\top}{\sqrt{d_k}}\right)V^k. \tag{7}$$

Multi-head attention is

$$\mathrm{MHA}(\mathbf{Z}) = \mathrm{Concat}(\mathrm{Attn}^1, \ldots, \mathrm{Attn}^H) W_O. \tag{8}$$

Each layer applies

$$\mathbf{Z}^{(\ell+1)} = \mathrm{LN}\Big(\mathbf{Z}^{(\ell)} + \mathrm{MHA}(\mathbf{Z}^{(\ell)})\Big), \tag{9}$$

followed by a positionwise feed-forward network

$$\mathrm{FFN}(\mathbf{u}) = \phi(\mathbf{u} W_1 + \mathbf{b}_1) W_2 + \mathbf{b}_2, \tag{10}$$

where $\phi(\cdot) = \max(0, \cdot)$. After $L$ layers, the encoder memory is

$$\mathbf{M} \in \mathbb{R}^{S \times d_{\mathrm{model}}}. \tag{11}$$

**Decoder masked self-attention** The teacher-forced decoder input $\tilde{\mathbf{y}}_{1:T} = (\mathbf{y}_0, \mathbf{y}_1, \ldots, \mathbf{y}_{T-1})$ is embedded analogously to obtain $\mathbf{U} \in \mathbb{R}^{T \times d_{\mathrm{model}}}$. Causality is enforced via mask

$$\mathbf{C}_{ij} = \begin{cases} 0, & j \le i, \\ -\infty, & j > i, \end{cases} \tag{12}$$

yielding

$$\mathrm{MaskedAttn}(Q, K, V) = \mathrm{softmax}\left(\frac{QK^\top}{\sqrt{d_k}} + \mathbf{C}\right) V. \tag{13}$$

**Encoder–decoder attention** Cross-attention uses

$$Q^k = \mathbf{U} W_Q^k, \quad K^k = \mathbf{M} W_K^k, \quad V^k = \mathbf{M} W_V^k, \tag{14}$$

allowing each decoder timestep to condition on $\mathbf{x}_{1:S}$.

**Autoregressive prediction** At timestep $t$, the decoder computes

$$\mathbf{h}_t^{\mathrm{dec}} = g(\hat{\mathbf{y}}_{1:t-1}, \mathbf{M}), \tag{15}$$

where $g(\cdot)$ denotes the stacked masked self-attention, cross-attention, and feed-forward transformations. The predicted coordinate is

$$\hat{\mathbf{y}}_t = W_{\mathrm{out}} \mathbf{h}_t^{\mathrm{dec}} + \mathbf{b}_{\mathrm{out}}, \quad W_{\mathrm{out}} \in \mathbb{R}^{2 \times d_{\mathrm{model}}}. \tag{16}$$
